# Conversion between parallel and antiparallel β-sheets in wild-type and Iowa mutant fibrils

**DOI:** 10.1101/220749

**Authors:** Wenhui Xi, Ulrich H.E. Hansmann

## Abstract

Using a variant of Hamilton-Replica-Exchange we study for wild type and Iowa mutant Aβ_40_ the conversion between fibrils with antiparallel β-sheets, and such with parallel β-sheets. We show that wild type and mutant form distinct salt bridges that in turn stabilize different fibril organizations. The conversion between the two fibril forms leads to the release of small aggregates that in the Iowa mutant may shift the equilibrium from fibrils to more toxic oligomers.

## INTRODUCTION

The pathogenesis of Alzheimer’s disease is correlated with the presence of insoluble amyloid fibrils,^1, 2^ assembled from 37-42 residue long Aβ peptides, that are the main component of the senile plaques found in AD’s patient brain.^3, 4^ The mature fibrils are formed by a nucleation process that involves assembly of small solvable oligomers now thought to be the main neurotoxic agent.^5, 6^ Monomers, oligomers and amyloid fibrils exist in an equilibrium of interchanging structures characterized by *polymorphism*.^7^ At least five different structures of Aβ_1-40_ amyloid fibrils, implicated in Alzheimer’s disease, have been determined by ssNMR^8-12^; and the differences in molecular structure are correlated with cell toxicity and disease progression.^12, 13^ One reason for the polymorphism are kinetic-traps and alternative pathways of aggregation,^14^ meaning that formation of fibrils and that of oligomers are competing processes. As some amyloid fibrils act as instruments for storing and releasing of hormones, one can speculate that under non-disease conditions, peptides and proteins are either in their functional form (that is folded or naturally disordered), or stored in fibrils.^15, 16^ On the other hand, under disease conditions, the off-pathway formation of oligomers dominates and leads to the onset of the disorders.

Environmental conditions such as inflammation-induced tissue-changes or metal ions^2^ may cause a shift toward disease conditions, but best studied are mutation-caused shifts. More than ten different Aβ mutants are known lead to familial forms of Alzheimer’s disease.^17^ While the disease-related structural changes caused by the mutation can be subtle, the situation is different for the Aβ_40_ Iowa mutant (D23N). Here, the mutation of residue 23 from an Aspartic Acid (D) to an Asparagine (N) leads to higher neurotoxicity and early onset of the disease^18^ that, as for the Italian mutant (E22K),^19^ is correlated with the presence of long-living meta-stable anti-parallel fibrils.^20^ While there is evidence that the wild type can also exist in an antiparallel manner,^20^ all experimental structures derived by solid-state-NMR for the wild type Aβ_40_ fibrils are build out of in-register parallel β-sheets.^10, 12^ On the other hand, Aβ_1-40_ Iowa mutant fibrils have either parallel or antiparallel β-sheets, see Figure 1.^21^ The latter are meta-stable, converting over time into fibrils with parallel structure, and have a slower seeding speed.^21^ Interestingly, Iowa mutant fibril fragments can seed wild-type Aβ_1-40_ monomers into fibrils with anti-parallel β-sheets.^22^ While parallel and antiparallel Iowa mutants differ little in cell toxicity,^21^ it appears likely that the different pathologies are related to the propensities with that the two forms are observed in wild type and Iowa mutant. Hence, for deriving therapeutic strategies it is important to know in detail the process by that the two forms convert into each other, and how the Iowa mutation shifts the equilibrium between parallel and antiparallel β-sheets toward antiparallel β-sheets.

**Fig. 1.**
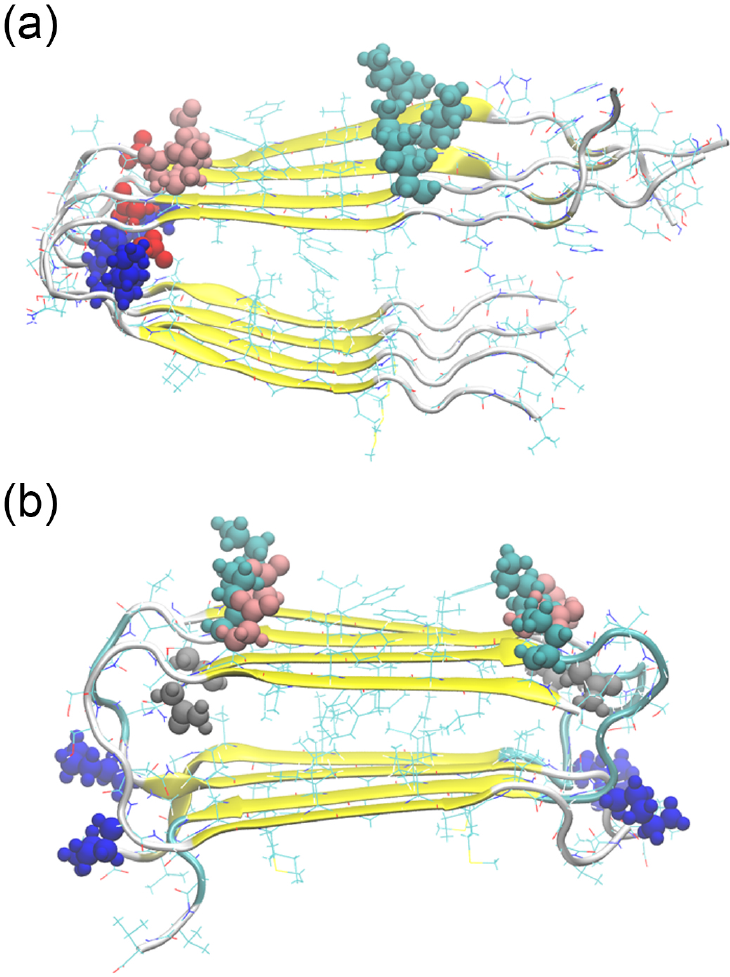
Parallel and antiparallel Aβ_40_ fibril models. All resolved Aβ_10_ wild type fibril models have parallel β-sheets (a), while the Iowa mutant D23N can also aggregate into fibrils with antiparallel sheets (b). Possible salt-bridge-forming side-chains of residues have been colored as follows: K16: cyan, K28: blue, E22: pink, D23 (wild type): red, N23 (Iowa mutant): gray.

It is taxing to probe the transition between parallel and antiparallel fibril structures in experiments or by computer simulations. Previous computational work comparing fibril-fragments of wild-type and Iowa mutant could not conclusively explain the conformational shift induced by D23N mutant.^23, 24^ Okamoto et al studied Iowa mutant dimers with both arrangements by molecular dynamics and ab initio fragment molecular orbital methods, focusing on the effect of the mutation on the parallel form.^25^ The stability of both structures was also investigated by us for wild type and Iowa mutant decamer with atomistic molecular dynamics simulations.^26^ We find that it is the alignment of hydrophobic interactions that makes for both wild type and Iowa mutant the in-register parallel form more stable than the antiparallel one. However, in agreement with recent experiments we find a higher stability for Iowa mutant antiparallel aggregates than for aggregates built of the wild type. MM-PBSA estimates of the free energy differences indicate that the transition from antiparallel to parallel forms, observed for the Iowa mutant, is costly. This suggests that this conversion is not a simple re-arrangement but rather involves the detachment of monomers from the antiparallel form and reassembly into the more stable parallel structure.

However, the transition pathway between parallel and antiparallel assembly is not known, and neither is the mechanism that enhances the propensity and life time of the antiparallel organization in the Iowa mutant. In this work, we try to answer these two questions through all-atom simulations that rely on a variant of the Hamilton Replica Exchange approach.^27, 28^ With this enhanced sampling method, we study the conversion between parallel and antiparallel fibril organizations of both wild type and Iowa mutant Aβ_15-40_ trimers. We find that in the Iowa mutant the mutation D23N increases the probability of forming an E22-K16 inter-chain salt-bridge which in turn enhances the formation of antiparallel fibril organization, while in the wild type the negatively charged residue D23 allows formations of intra-chain salt bridge D23-K28 that stabilizes the parallel form. The transition between the two fibril organizations requires partial dissolution of the fibrils which into smaller aggregates which in turn increases the frequency of potentially cytotoxic oligomers. Hence, we conjecture that the early onset and more severe symptoms of Alzheimer’s disease in patients with the Iowa mutant is not directly caused by the higher frequency of antiparallel fibrils, but instead results from the conversion of the metastable antiparallel fibrils into parallel ones, which shifts the equilibrium from less toxic fibrils toward more toxic oligomers.

## MATERIALS AND METHODS

### System construction

In order to research the differences between wild type and Iowa mutant fibrils, we perform Hamilton replica exchange (H-REMD) simulations^27,28^ of wild type and Iowa mutant Aβ_15-40_ trimer. Trimers are chosen because they are the minimal system size to model a fibril architecture (with an internal chain and chains at the boundaries) that is sufficiently stable but at the same time small enough to allow for large-enough all-atom simulations. For this purpose, we first generate trimer fibril fragments using as starting point for the parallel organization the wild-type Aβ_9–40_ fibril model (PDB code: 2LMO), and for the antiparallel organizations the antiparallel form of D23N Aβ_15–40_ fibril models (PDB code: 2LNQ). To have the same length we remove the residues 9 to 14 of the wild type model. For the same reason we use only one of the two layers in the 2LMO wild type fibril model. The chains are capped with acetyl and amide groups at the terminals to compensate for the extra charge group introduced by the residue deletion. The parallel Iowa mutant fibril fragment is now build by altering the residue D23 (an Aspartic Acid) in the wild type model into an Asparagine (23N) using the program UCSF Chimera,^29^ and regard the resulting structure as a parallel fibril model for the Iowa mutant. In a similar way, an antiparallel wild type fibril fragment is generated by replacing residue N23 into D23 in the Iowa mutant fibril model. The resulting four trimers are minimized and relaxed in a 1 ns long molecular dynamics run at 275K.

### Simulation Set-up

Our approach relies on introducing a potential energy made out of three terms:

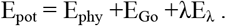

The first term is the “physical” energy E_phy_ of the amyloid and the surrounding environment described by an *all-atom* model. The second term E_Go_ describes our system by suitable *Go* model that bias toward distinct configurations. The degree of bias by the Go-model varies with replica since both models are coupled by a *system-specific penalty* term E_λ_, proposed in Ref.^30^ and tested extensively in a different context in Ref.^31^, that measures their similarity. The strength by that the two models are coupled is set by the replica-specific parameter λ.

Exchanging replicas with a probability

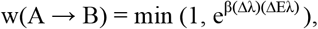

where Δλ = λ_B_ – λ_A_ and ΔE_λ_ = E_λ_(B) – E_λ_(A), configurations will walk along a ladder of replicas that mix to various degrees contributions from a Go-model with that from a physical force field. Our set-up introduces on one end of the ladder a bias for parallel fibrils and on the other side for anti-parallel forms. However, measurements are made solely at a “physical” replica which has no contributions from the Go-model (λ = 0). Such “feeding” of physical systems by Go-models has been proposed by us^32, 33^ and others^31, 34^ as a way to avoid the intrinsic bias in Go-model simulations. Note that our and the earlier approaches suffer often from the problem that the acceptance rate becomes vanishingly small for exchanges with the λ = 0 replica if not a large number of replicas is concentrated around it. We have developed a technique, called Replica-Exchange-with-Tunneling (RET) to overcome this problem^35^, but due to the small system size was use of RET not necessary in the present study. However, use of RET would be required if we wanted to study larger oligomers.

The above algorithm has been implemented by us into the often used software package GROMACS 4.6.5.^36^ We use the CHARMM27 force field with CMAP corrections^37^ in the “physical” model as this allows us to compare results with our previous work.^26^ In order to speed up simulation, we estimate protein-solvent interactions by a generalized-Born implicit solvent model^38^. The biasing Go-model parameters are generated by the SOMG-server^39^ using as input the above generated reference structures. In each Hamilton Replica Exchange run, 16 replicas are used with λ = (0.015, 0.0135, 0.012, 0.01, 0.0075, 0.005, 0.002, 0.0, 0.0, 0.002, 0.005, 0.0075, 0.01, 0.012, 0.0135, 0.015). Noticing that the replica 7 and 8 is not biased with λ=0. The thermostat temperature are controlled by the stochastic v-rescaling method^40^ and change from 310K to 310.15K with step of 0.01K. The cutoff is set to 1.5nm for both van der Waals (vdW) and electrostatic interactions. Bond lengths of all hydrogen atoms are constrained with the LINCS algorithm^41^ in the physical model. Exchanges are tried every 20ps and the integration time step is 2 fs with leapfrog algorithm. Snapshots of configurations are also saved each 20ps. Each replica trajectory undergoes 200ns and only the last 100ns data are used for analysis.

### Analysis

Most of our analysis relies on the GROMACS tools suite. The free energy landscape of wild type and mutant fibril trimers are calculated by reweighing from replica with λ≠0 to λ=0 where the physical model is not biased by a Go-term to either parallel or antiparallel fibril organization, for more technical details see our previous work.^42^ Comparing re-weighted landscapes for different time sections we checked for convergence of our simulation, and found that we had to omit the first 100ns to ensure that our systems are in equilibrium. Hence, our results are based on data collected in the time interval of 100-200ns. Salt-bridges are defined by the distance between the mass center of COO- and NH3+ groups in the side chains of amino acid being smaller than 0.45nm. Note that the artificial dynamics in our sampling method does not allow us to extract dynamics in a naïve way. Instead, we follow our previous work^42^ and employ a variant of Markov state models^43^ to classify all configuration and to probe the transitions between kinetically relevant states. For his purpose, we have combined all 16 trajectories, and selected configuration every 40ps for the last 100 ns. Using the Gromacs routine *g_rms* we then calculated the RMSD matrix for these 40,000 configurations. The resulting matrix allowed us to use the Ward method for hierarchical clustering in R language^44^ to group the sampled configurations. Using the procedure described in Ref.^42^, we found a number of 20 clusters optimal for our analysis. The weight of each cluster is the sum of the weights of its members, but note, that the weight of a member configuration has to be calculated by re-weighting if this configuration was not sampled at a replica with λ=0.

## RESULTS

### Free energy landscape of wild type and Iowa mutant

Our Hamilton-Replica-Exchange realizes a random walk in λ-space between replicas where a Go-model strongly biases the physical model toward fibril-fragments with parallel β-sheets, and such on the other side of the ladder where the Go-model biases toward the antiparallel organization. The net-effect is sampling of a larger number of *independent* configurations at the central replicas where λ=0, i.e., where there is no bias by any Go-model. The available statistics can be further increased by reweighting data from the other replicas toward λ=0. The enhanced sampling allows us to derive the free energy landscape of the two systems, shown together with typical configurations in Figure 2a for wild type and in Figure 2b for the Iowa mutant. The landscape is in both cases projected on the root-mean-square-deviation (RMSD) with either the antiparallel fibril organization (x-axis) or the parallel form (y-axis), and free energies are given in units of RT. For both wild type and Iowa mutant fibril, we find that parallel fibril organizations have the lowest free energy. In order to compare the two systems, we have shifted the free energies such that the global minimum in each system takes a value of zero. Comparing wild type and Iowa mutant we find that the antiparallel fibril fragment has a 2.9 RT higher free energy for the wild type, while the corresponding value for the Iowa mutant is only 2.6 RT. While this difference is consistent with our previous simulations^26^ and experimental work,^45^ its value is too small to explain the different life times of antiparallel fibrils in wild type and mutant: experimentally, antiparallel and parallel fibril of Iowa mutants form at the same time and coexist for about 100 hours before the antiparallel form gradually dissolves.^21^ These different life times are rather due to the bottlenecks in the landscape that are much more pronounced for the wild type for the mutant.

**Fig. 2.**
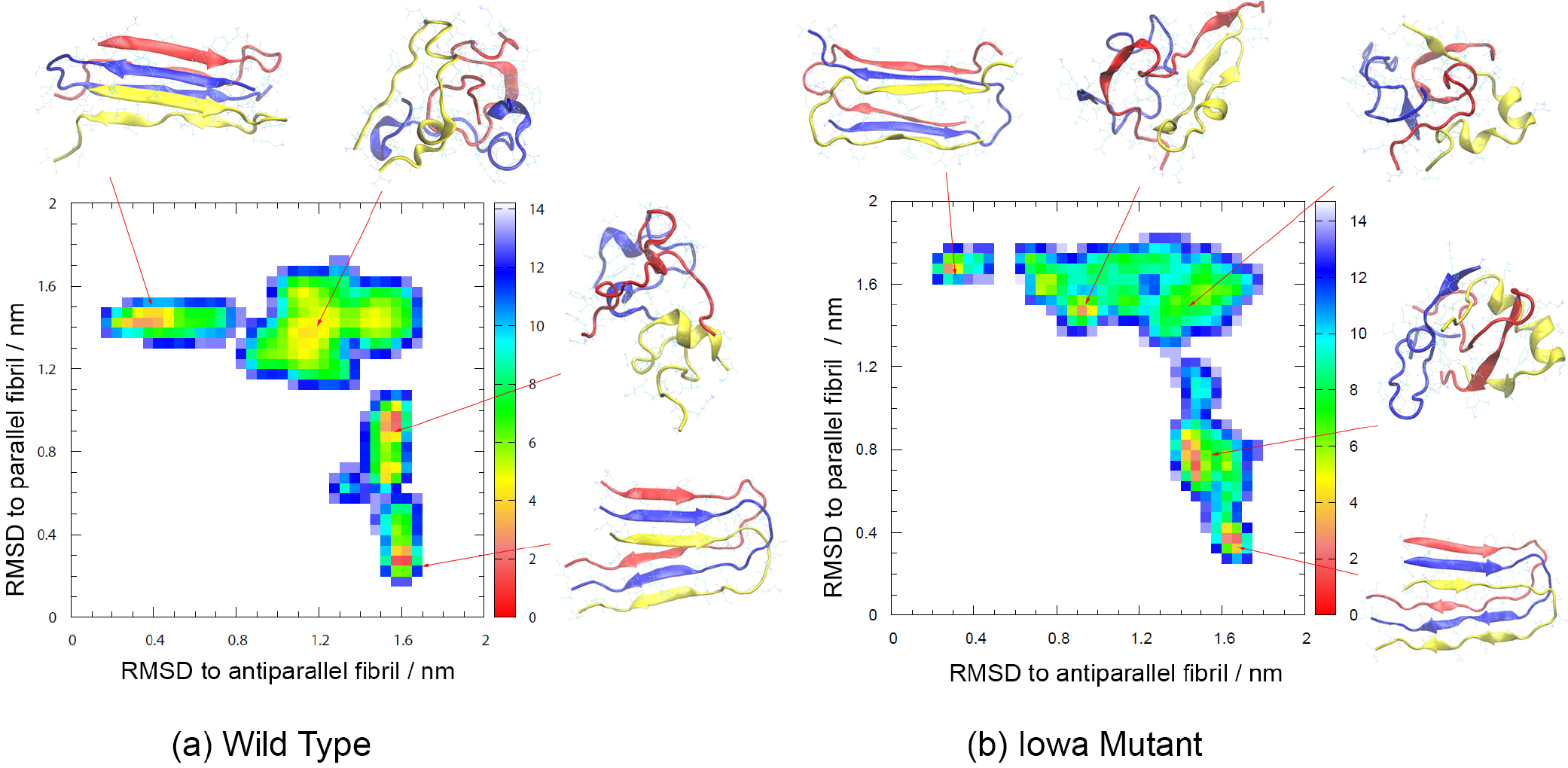
Free energy landscape of wild type(a) and Iowa mutant Aβ_15-40_ trimers projected on the root-mean-square-deviation (RMSD) with either parallel or antiparallel fibril organization. Free energy values are given in RT and shifted such that in each of the two systems the minimum value is zero. Representative structures for the various basins are also shown.

Residue-contact maps offer another way to deduce mechanism and packing pattern. As an example, we show in Fig. 3 the backbone contact maps for both Iowa mutant and wild type. The contact maps are divided into intra-chain (Fig. 3 a) and c)) contacts and inter-chain contacts (Fig. 3 b) and d)). We observe in both maps a higher content of antiparallel β-sheets for the mutant than for the wild type. Especially, the intra-chain contacts in the region V32- I34 are formed only for the Iowa mutant. Similarly, the inter chain contacts between residues in the segment L17 and E22, located in the central hydrophobic core region of Aβ peptides, are formed only for the Iowa mutant, while in the wild type such inter-chain antiparallel β-sheet contacts are seen only in the region I32-V36.

**Fig. 3.**
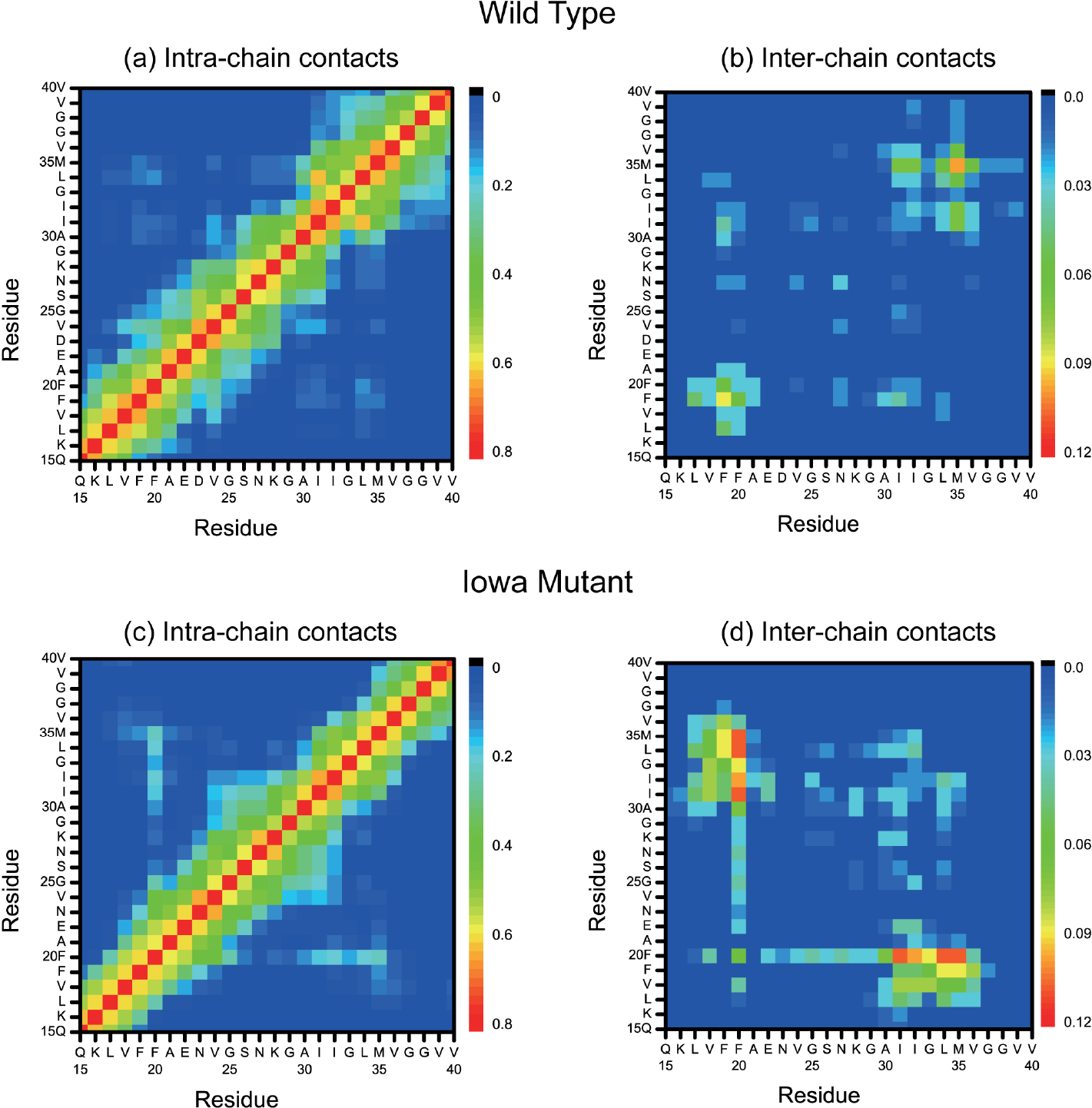
Intra-chain and inter-chain backbone contacts in wild type (a, b) and Iowa mutant (c, d) fibril fragments.

Of special importance are the contacts formed by residue 23. In the wild type this is a negatively charged Aspartic Acid (D) forming in the parallel fibril structure a salt bridge D23-K28 with the positively charged Lysine (K) at position 28, whose importance for Aβ_40_ fibril aggregation is well-known.^10, 21, 46^ In the Iowa mutant, the negatively charged D23 is replaced by a neutral (polar) Asparagine (N23), i.e., residue 23 can no longer form a salt bridge with residue K28. However, while in the wild type the salt bridge D23-K28 forces the side-chain of residue E22 (a negatively charged Glutamic Acid) to the outside of the fibril,^10^ this constraint does not exist in the Iowa mutant, and residue E22 can now form an inter-chain salt-bridge with the positively charged residue K16, when the Iowa mutant chains are in the antiparallel fibril form.^45^ As wild type and mutant differ in what salt bridges they can form, we have measured the frequencies of the different possible inter-chain/intra-chain salt-bridges, and show these frequencies in Figure 4. As expected, the intra-chain D23-K28 salt bridge is the one with highest probability for the wild type; and while residue K28 can also form inter-chain or intra-chain salt-bridges with residue E22, these appear with much lower frequency. On the other hand, the D23-K28 salt bridge cannot be formed in the Iowa mutant trimer. Instead, the inter-chain K16-E22 salt-bridge is now seen with high frequency, and to a smaller degree E22-K28 salt-bridges. Hence, in the wild type, the residues E22 and D23 compete in forming contact with residues K28 or K16 which makes it difficult to form the K16-E22 salt-bridge that is essential in antiparallel assemblies. However, the mutant D23N cannot form the salt-bridge D23-K28 that stabilizes parallel fibrils, and without competition from D23, the inter-chain K16-E22 salt-bridge appears with high frequency stabilizing therefore the antiparallel structure.

**Fig. 4.**
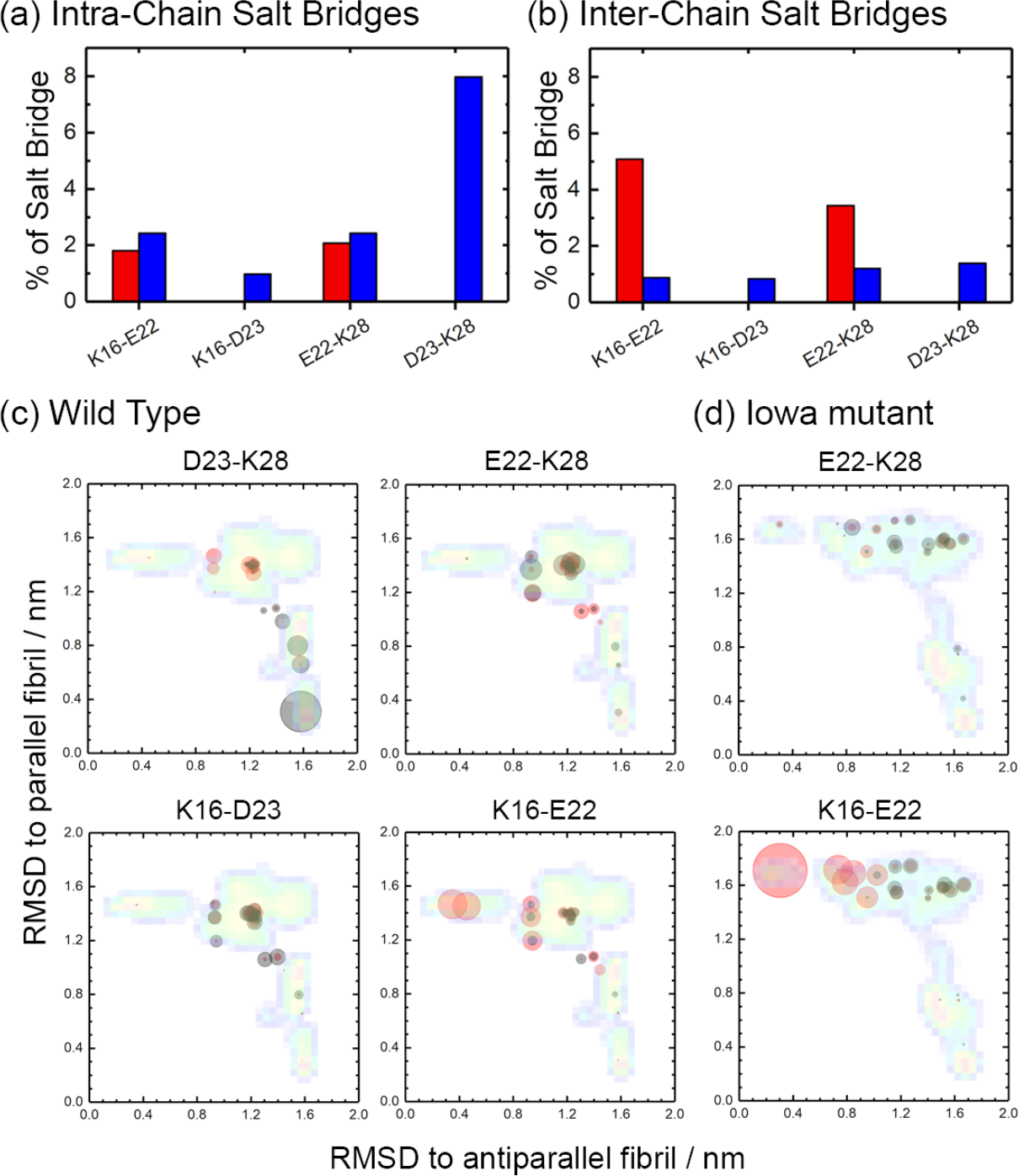
Frequency of possible intra-chain (a) and inter-chain (b) salt-bridges involving positively charged residues K16, K28, and negatively charged residues E22 and D23, as measured in our simulations of wild type (red) and Iowa mutant (blue) Aβ_15-40_ trimers. The average frequency of the various possible salt bridges for wild type (c) and Iowa mutant (d) is also shown as an overlay on the free energy landscape of Fig. 2. Frequencies of inter-chain salt bridges are represented by red circles, and the ones of intra-chain salt bridges by black circles.

A more detailed view is given in Figure 4 (c) and (d) where we show the average frequency of each salt bridge, projected on the same coordinates as the free energy landscape of Figure 2. For this purpose, we have divided the sampled configurations into 20 clusters as described in the Method section, and calculated the weighted average frequency for each cluster. Each cluster is marked by a circle, with the center of the circle given by the central member configuration of the cluster, and its radius given in units of pixel by 40 times the average weighted frequency for a certain salt bridge. Comparing wild type and Iowa mutant, we see that salt bridges involving residue E22 appear almost exclusively in configurations with antiparallel sheets or in the transition region, but never in parallel fibrils. With significant probability one finds only inter-chain K16-E22 salt bridges, and these mostly for antiparallel fibrils. As already seen earlier, the frequency of this salt bridge is much higher in the mutant than in the wild type, i.e., the antiparallel fibril organization has in the Iowa mutant an additional stabilization by this salt bridge that is missing in the wild type. On the other hand, salt bridges involving residue D23, which are only possible in the wild type, are never observed in fibrils with antiparallel organization. Instead, they are overwhelmingly located in parallel fibrils as intra-chain salt bridge D23–K28. Inter-chain salt bridges and such between D23-K16 exist, but only with low frequency and in the transition region. Hence, the wild type is not only missing the stabilization of antiparallel fibrils by an K16-E22 salt bridge seen in Iowa mutant fibrils, but in addition, parallel fibrils have an extra stabilization by D23-K28 intra-chain salt bridges. These different stabilizing salt bridges explain why meta stable antiparallel fibrils can exist for the Iowa mutant; but almost never seen for the wild type, where this organization is energetically weaker and on the other side the parallel fibrils more stable.. Note also that in the wild type the transition state region is more populated by an ensemble of diverse salt-bridge stabilized configurations than seen for the Iowa mutant which may make it kinetically easier to go in the wild type into the parallel fibril form, However, when seeded with antiparallel fibril fragments wild type chains can form inter-chain K16-E22 salt bridges with the seed and assemble into metastable antiparallel fibrils^22^, While the relative stability of these fibrils compared to antiparallel Iowa mutant fibrils is not known, we would expect the antiparallel wild type fibrils to decay much faster into the parallel form than the Iowa mutant fibrils as residue D23 interferes with E22 and an intra-chain salt bridge D23-K28 is more favorable in the wild type.

Both antiparallel and parallel β-sheet Iowa mutant fibrils are cytotoxic at concentrations of μM level. However, their toxicity differs little between themselves and from the (parallel) wild type Aβ_40_ fibrils, but is much smaller than that of wild type Aβ_40_ oligomers at similar concentrations.^47^ This suggests that the different pathologies of wild type and Iowa mutant do not result directly from the propensities of the two fibril architectures in wild type and mutant, but are rather related to the frequency of transient oligomers formed in the transition between antiparallel and parallel fibrils. In order to test the plausibility of this hypothesis and learn more about the conversion process, we show in Figure 5 (a) and (c) the frequency of inter-chain hydrogen bonds seen in either parallel to antiparallel fibril organization. The frequencies are again calculated for the 20 clusters into which we have dived the sampled configurations. Each cluster is again represented by a circle, with the radius indicating the frequency of hydrogen bonds, and the clusters drawn as an overlay onto the free energy landscape of Figure 2. For both wild type and Iowa mutant we see large circles in the region of either parallel or antiparallel fibrils, marking the large number of inter-chain hydrogen bonds keeping the trimers together in the respective fibril organization. On the other hand, the circles are much smaller, or not seen, in the transition region between the two forms, indicating that the fibril fragments have dissolved into an ensemble of single chains. Hence, we conclude that the conversion between antiparallel and parallel fibrils requires dissolution of the fibril and release of monomers. We conjecture that in a cell these monomers will not always reassemble immediately into the other fibril organization, but may also mis-assemble into the oligomers that are the main cytotoxic agent.

**Fig. 5.**
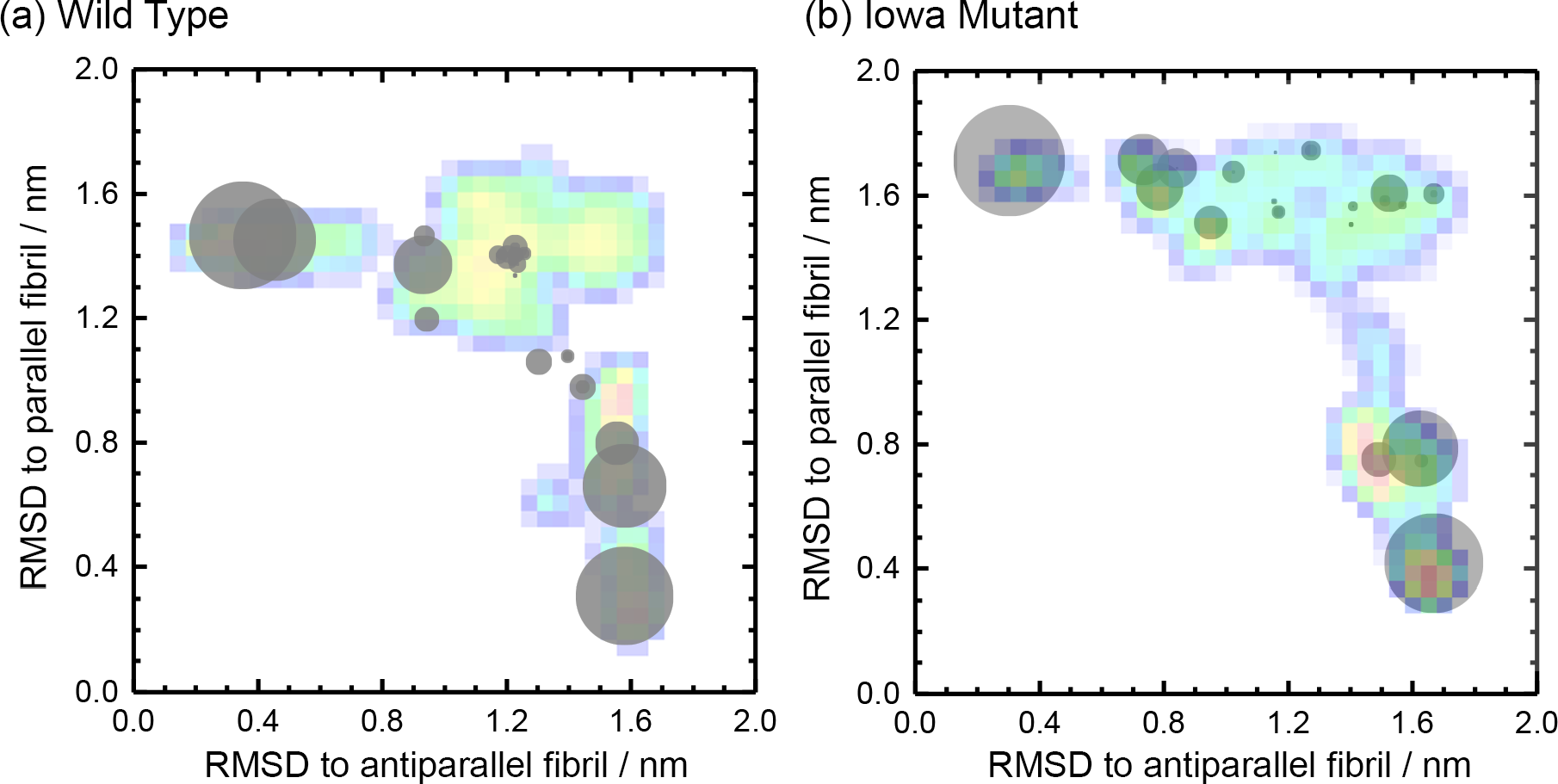
The frequency of inter-chain hydrogen bonds seen in either parallel or antiparallel fibril organization for wild type (a) and Iowa mutant (b) as an overlay on the free energy landscape of Fig. 2. The radius of circle is given by the frequency.

## DISCUSSION AND CONCLUSION

Using a variant of Hamilton replica exchange molecular dynamics, we have studied wild-type and Iowa mutant trimmers of Aβ_15–40_ peptides. Our enhanced sampling method allowed us to sample the free energy landscape of the two systems and the conversion between fibrils with antiparallel β-sheets, only seen as long-living meta-stable state in the Iowa mutant, and such with parallel β-sheets, seen in both wild type and mutant. We find that in the wild type the charged residue D23 allows formation of an intra-chain salt bridge D23-K28 that stabilizes parallel fibrils. In the Iowa mutant this salt bridge cannot be formed as the mutation D23N replaces a charged residue by a polar one. Instead, it becomes possible in the Iowa mutant to form an inter-chain salt bridge K16-E22 that stabilizes and enhances formation of the antiparallel fibril organization, which becomes now sufficiently stable to exist as long-living metastable state in the Iowa mutant. In our simulations requires the conversion between antiparallel and parallel fibril organization decay of the respective fibril fragments and release of monomers. We speculate that in the cell these monomers, or other small aggregates released in the conversion, do not always reassemble immediately into the other fibril organization. Instead, they may serve as a reservoir for formation of the toxic oligomers that are thought to be the main agent causing the disease symptoms in Alzheimer’s disease. Hence, the early onset and more severe symptoms of Alzheimer’s disease in patients with the Iowa mutant may not be directly related to the higher frequency of metastable antiparallel fibrils, but because their conversion into fibrils with parallel β-sheets shifts the equilibrium from less toxic fibrils toward more toxic oligomers.

## ACKNOWLEDGMENTS

The simulations in this work were done using the SCHOONER cluster of the University of Oklahoma and XSEDE resources allocated under grant MCB160005 of the National Science Foundation. We acknowledge financial support from the National Institutes of Health under grant GM120578.

